# NHBA is processed by kallikrein from human saliva

**DOI:** 10.1101/395988

**Authors:** Elisa Pantano, Sara Marchi, Massimiliano Biagini, Martina Di Fede, Vincenzo Nardi-Dei, Silvia Rossi Paccani, Mariagrazia Pizza, Elena Cartocci

## Abstract

Neisserial Heparin Binding Antigen (NHBA) is a surface-exposed lipoprotein and a component of the Bexsero vaccine. NHBA is characterized by the presence of a highly conserved Arg-rich region involved in binding to heparin and heparin sulphate proteoglycans present on the surface of host epithelial cells, suggesting a possible role of NHBA during *N. meningitidis* colonization. NHBA has been shown to be cleaved by the bacterial NalP protein, a meningococcal protease and by human lactoferrin (hLF), a host protease present in different body fluids (saliva, breast milk and serum). Cleavage occurs upstream or downstream the Arg-rich region. Since the human nasopharynx is the only known reservoir of infection, we further investigated the susceptibility of NHBA to human proteases present in the saliva to assess whether proteolytic cleavage could happen during the initial steps of colonization. Here we show that human saliva proteolytically cleaves NHBA; and identified human kallikrein 1 (KLK1) as the main protease responsible for this cleavage. Kallikrein is an important enzyme present in blood plasma, lymph, urine, saliva, pancreatic juices, and other body fluids that catalyze the proteolysis of several human proteins. We report the *in vitro* characterization of NHBA cleavage by kallikrein; the identification of the cleavage in the recombinant NHBA protein and, on the native protein, when expressed on live bacteria. Overall, this findings provide new insights on NHBA as target of host proteases, highlights a potential role of NHBA in the *Neisseria meningitidis* nasopharyngeal colonization, and of kallikrein as a defensive agent against meningococcal infection.

## Introduction

*Neisseria meningitidis* is a pathogenic, encapsulated, aerobic gram-negative diplococcus, member of the *Neisseriae* family. It is an obligate commensal in man and colonizes the nasopharyngeal mucosa without affecting the host, a phenomenon known as carriage recognized since 1890 **[1].** In nonepidemic settings, approximately 10% of healthy individuals at any time carry *Neisseria meningitidis* in the oropharyngeal tract. Transmission of bacteria occurs via direct contact or through dispersion of respiratory droplets from a healthy carrier or an infected person to a susceptible individual. Although often protected by a polysaccharide capsule, meningococci are particularly sensitive to desiccation; thus, spread from one individual to another necessitates close contact **[4].** In fact, closed or semi-closed settings, such as residential schools and military recruit camps, facilitate meningococcal transmission resulting in a dramatic increase of carriage rate, which may approach 100%.

Occasionally, shortly after the onset of colonization, *N. meningitidis* penetrates the mucosal membrane and enters the bloodstream, causing various forms of disease [5]. The most common clinical presentations of invasive infections include meningitis and severe sepsis with an often fatal outcome; other diseases, such as septic arthritis, pneumonia, purulent pericarditis, conjunctivitis, otitis, sinusitis and urethritis, occur more rarely **[5]**.

Meningococcal serogroups B and C are responsible for the majority of meningitidis cases in Europe, Y serogroup is relevant in the United States **[8]**, whereas African epidemics are mainly attributable to the A strain. While effective vaccines based on capsular polysaccharides against serogroups A, C, W and Y are avialble, different strategies have been pursued for developing a vaccine against the serogroup B being its capsular polysaccharide structurally identical to a component of the human fetal neural cell-adhesion molecule **[24]**. In 2000, genome mining and subsequent reverse vaccinology analysis brought to the discovery of three main protein antigens. These proteins, present as single antigen or as fusion protein, were formulated with the Outer Membrane Vesicles from a New Zealand outbreak strain (NZ98/254) shown to be protective in a immunization campaign in New Zealand **[6]**. Nowadays, Bexsero, is a multicomponent vaccine against *Neisseria meningitidis* serogroup B **[20]**.

Neisseral Heparin Binding Antigen (NHBA), previously named as GNA2132 (Genome-derived Neisseria Antigen 2132), is one of the three main protein antigens of the Bexsero vaccine. NHBA is a surfaced-exposed lipoprotein, its *nhba gene* is ubiquitous in meningococcal strains of all different serogroups and it has also been found in several other *Neisseria* species, including *N*. *lactamica*, *N. polysaccharea* and *N. Flavescens* **[7].** Recent studies on NHBA have shown that this protein is implicated in different steps of meningococcal pathogenesis, including bacterial adhesion to epithelial cells **[16]**, biofilm formation **[25]**, bacterial survival in the blood, and vascular leakage **[15]**. It is also able to bind heparin *in vitro* through an Arginine-rich region **[7]** and this ability may point to a role of NHBA in the protection of unencapsulated meningococci against complement **[15]**.

The primary amino-acid sequence of NHBA, from strain 2996, comprises approximately 450 residues and its structure can be divided into two main domains: an N-terminal region (residues 1 to ∼230), marked as an intrinsically unfolded region, and a C-terminal region, whose structure, organized in β-barrel (residues 305-426) with high thermal stability, has been solved recently by NMR spectroscopy and X ray diffraction **[20] [21]**. A highly conserved Arginine-rich motif, responsible for protein binding to heparin and heparin sulfate proteoglycans **[15] [7]**, (residues 235– 245) is located between the N-and C-terminal regions **[20] [21]**. It has been reported **[7]** that NHBA can be processed upstream or downstream of the Arg-rich region by the bacterial NalP protease and by human lactoferrin (hLF), respectively. Since hLF is a protease present in different body fluids, including saliva, we hypothesized that NHBA protein could be cleaved by proteases contained in the saliva during the initial step of *N. meningitis* colonization of the nasopharyngeal mucosa. Here we show that NHBA is proteolysed during incubation with saliva and identified allikrein 1 as responsible for the cleavage. The NHBA fragments generated by kallikrein cleavage were characterized by mass spectrometry and shown to be identical to the NHBA fragments generated by human lactoferrin proteolytic cleavage.

Finally, we show that the native NHBA surface exposed on live bacterial surface is cleaved by saliva and hKLK1 and that the cleavage results in the release of the C-fragment in the culture supernatant.

## Materials and Methods

### SDS-PAGE and Western Blot analysis

Proteins were separated by SDS-PAGE electrophoresis using 4-12 % or 12 % polyacrylamide NuPAGE Bis-Tris Precast Gels (Invitrogen). For SDS-PAGE analysis, gels were stained with SimplyBlue Safe Stain (Invitrogen). For Western blot (WB) analysis, 1 ml of culture medium supernatant was recovered, filtered 0.22 μm and precipitated in 10% (v/v) trichloroacetic acid. Protein pellets were solubilized in 50 μl of Novex LDS Sample Buffer 1X (Thermo Fisher) containing Tris Base 0.1 M and NuPage Sample reducing agent 1X (Thermo Fisher). Precipitated proteins were run on SDS-Page and the gel was transferred onto nitrocellulose membranes. Western blots were performed according to standard procedures. NHBA full-length protein and its C-terminal fragments were identified with polyclonal mouse antisera raised against the recombinant NHBA full-length protein (working dilution 1:1000) or against the recombinant C2 fragment (working dilution 1:1000), respectively. An anti-mouse antiserum conjugated to horseradish peroxidase (32230 Thermo Fisher) was used as secondary antibody. Bands were visualized with Super Signal West Pico Chemiluminescent Substrate (Thermo Fisher) following the manufacturer’s instructions.

### Expression and purification of recombinant NHBA proteins

Recombinant wt and Δ-Arg NHBA protein was expressed in *E. coli* BL21 (DE3) strain by using EnPresso B growth kit (BioSilta) supplemented with 100 μg/ml ampicillin. Bacteria were grown at 30°C for 16 h, and protein expression was induced by the addition of 1 mM isopropyl β-D-1-thiogalactopyranoside (IPTG) (Sigma) at 25°C for 24 h. The soluble proteins were extracted by sonication in 50 mM NaH_2_PO_4_, 300 mM NaCl, 10 mM imidazole, pH=8, supplemented with protease inhibitors (cOmplete™, EDTA-free Protease Inhibitor Cocktail, Roche), followed by centrifugation to remove cell debris. Recombinant proteins were purified from 0.22 μm filtered supernatant by affinity chromatography with HisTrap HP column (GE Healthcare) using an AktaPurifier System (GE Healtcare) following the recommended procedure. After washing, elution was performed by gradient step from 0 to 250 mM imidazole in 20 CV. Eluted sample was dialyzed at 4 °C overnight against 10 mM NaH2PO4, pH=7, buffer using Snake Skin Dialysis Tubing, 10,000 MWCO, 22 mm (Thermo Scientific), and a second step of purification was performed to obtain a high pure product. WT NHBA dialyzed sample was load on HiTrap heparin HP column (GE Healthcare) using an AktaPurifier System (GE Healtcare) following the recommended procedure. After washing, elution wasperformed by gradient step from 0 to 500 mM NaCl. Δ-arg NHBA dialyzed sample was load on HiTrap Q HP column (GE Healthcare) using an AktaPurifier System (GE Healtcare) following the recommended procedure. After washing, elution was performed by gradient step from 0 to 500 mM NaCl. Fot both sampls, to exchange the buffer, eluted samples were dialyzed at 4 °C overnight against PBS using Snake Skin Dialysis Tubing, 10,000 MWCO, 22 mm (Thermo Scientific). Protein content was quantified using the BCA Kit (Themo Fischer Scientific). Purity was checked by SDS-PAGE analysis, as well as by SEC-UPLC loading the sample on BEH200 4.6×300mm column (Waters). Size exclusion chromatography was performed using 10 mM NaH_2_PO_4_, 400 mM (NH_4_)_2_SO_4_, pH=6, buffer. Endotoxin content was quantified by LAL assay using ENDOSAFE instrument (Charles River).

### NHBA in vitro cleavage by saliva and kallikrein1

NHBA recombinant proteins, obtained as previously described, were incubated with saliva (Human saliva lot BRH1325364 from Seralab) in a ratio of 1 ml of saliva per mg of recombinant proteins, or with recombinant human kallikrein1 (2337-SE R&D System) or with kallikrein1 purified from human plasma (K2638 Sigma Aldrich) at different ratio, from 1:500 to 1:500000. Samples were incubated at 37°C overnight; then they were centrifuged for 1 minute and analysed on SDS-PAGE.

### Proteomic analysis of saliva

For proteomic analysis, human saliva (Human saliva lot BRH1325364 from Seralab) was fractionated by anion exchange chromatography. Dialyzed saliva has been loaded on Column Mini Q PE 4.6/50 GE Healthcare (0.8 ml CV), with a flow of 0.2 ml/min on a Acquity-HPLC system (Waters) previously equilibrated with buffer 25 mM Tris pH 8.0.

Elution was performedwith a flow of 1 ml/min in a gradient mode: from 250 mM Tris, pH 8, to 25 mM Tris HCl 0,5 M NaCl, pH 8 in 20 CV and proteins were collected in fractions of 0,5 ml volume. NHBA cleavage activity of each fractions were evaluated incubating NHBA at a concentration of 1 mg/ml with an equal volume of each fraction as previously described. Protein fractions positive for proteolytic activity on NHBA protein were pooled and precipitated in 10% (v/v) trichloroacetic acid and 0.04% (w/v) sodium dehoxicholate. Protein pellets were solubilized in 50 μl di 0.1% (w/v) Rapigest^™^ (Waters, MA, USA) and 1mM DTT and 50 mM ammonium bicarbonate, boiled at 100°C for 10 min. After cooling down, 1μg of LysC/trypsin mix (Promega) was added and the reaction was performed overnight. Digestions were stopped with 0.1% final formic acid, desalted using OASIS HLB cartridges (Waters) as described by the manufacturer, dried in a Centrivap Concentrator (Labconco) and resuspended in 100 μl of 3% (v/v) acetonitrile (ACNcan) and 0.1% (v/v) formic acid. An Acquity HPLC instrument (Waters) was coupled on-line to a Q Exactive Plus (Thermo Fisher Scientific) with an electrospray ion source (Thermo Fisher Scientific). The peptide mixture (10 μl) was loaded onto a C18-reversed phase column Acquity UPLC peptide CSH C18 130Å, 1.7μm 1 x 150 mm and separated with a linear gradient of 28–85% buffer B (0.1% (v/v) formic acid in ACN) at a flow rate of 50 μL/min and 50 °C. MS data was acquired in positive mode using a data-dependent acquisition (DDA) dynamically choosing the five most abundant precursor ions from the survey scan (300–1600 *m/z*) at 70,000 resolution for HCD fragmentation. Automatic Gain Control (AGC) was set at 3E+6. For MS/MS acquisition, the isolation of precursors was performed with a 3 *m/z* window and MS/MS scans were acquired at a resolution of 17,500 at 200 *m/z* with normalized collision energy of 26 eV. The mass spectrometric raw data were analyzed with the PEAKS software ver. 8 (Bioinformatics Solutions Inc., ON, Canada) for de novo sequencing, database matching and identification. Peptide scoring for identification was based on a database search with an initial allowed mass deviation of the precursor ion of up to 15 ppm. The allowed fragment mass deviation was 0.05 Da. Protein identification from MS/MS spectra was performed against NCBInr *Homo sapiens* (Human) protein database (112,970,924 protein entries; 41,399,473,309 residues) combined with common contaminants (human keratins and autoproteolytic fragments of trypsin) with a FDR set at 0.1%. Enzyme specificity was set as C-terminal to Arg and Lys, with a maximum of four missed cleavages. N-terminal pyroGlu, Met oxidation and Gln/Asn deamidation were set as variable modifications.

### Purification of NHBA fragments after cleavage

NHBA recombinant protein was digested with saliva and recombinant KLK1 as previously described and the obtained digested fragments were separated on His Spin Trap devices (GE Healtcare) following the costumer procedure. The His Spin Trap was equilibrated with 50 mM NaH_2_PO_4_ pH8, 300 mM NaCl, NHBA cleaved recombinant protein was loaded on the devices and after washing C-fragments waseluted with 50 mM NaH_2_PO_4_ pH8, 300 mM NaCl, 500 mM Imidazol.

### Mass Spectometry analysis of the cleaved purified C-term fragments

The C fragment recovered from His Spin Trap elution fractionwas analysed by Mass Spectrometry. The acidified protein solutions were loaded onto a Protein Microtrap cartridge (from 60 to 100 pmols), desalted for 2 min with 0.1% (v/v) formic acid at a flow rate of 200 ml/min and eluted directly into the mass spectrometer using a step gradient of acetonitrile (55% (v/v) acetonitrile, 0.1% (v/v) formic acid). Spectra were acquired in positive mode on a SynaptG2 HDMS mass spectrometer (Waters) equipped with a Z-spray ESI source. The quadrupole profile was optimized to ensure the best transmission of all ions generated during the ionization process. Mass spectra were smoothed, centroided and deconvoluted using MassLynx vers. 4.1 (Waters).

### Bacterial strains and growth conditions

*N. meningitidis* wt strain MC58 **[22]** and the ΔNalP and ΔNHBA MC58 strains **[7]** were cultured on GC agar plates (Difco) at 37 °C plus 5% CO_2_. Overnight colonies grown on GC agar plate were used to start a culture in 25ml of GC medium (Difco) in flask: OD_600nm_ ∼ 0.1 was used as start culture density. Liquid culture was grown at 32 °C plus 5% CO_2_, under aerobic condition (180 rpm) until OD_600nm_ ∼ 0.4 and then used for experimentation. Bacteria were incubated with saliva (Human saliva lot BRH1325364 from Seralab), human plasma-purified KLK1 (K2638 Sigma Aldrich) or liquid GC medium, as negative control, for 30 minutes. Culture medium supernatant was recovered and used for Western blot analysis; while bacteria were used for FACS analysis.

### Flow Cytometry analysis

Bacteria, obtained as previously described, were washed and incubated for 1 hour, at room temperature, with polyclonal mouse antiserum raised against NHBA C-terminal fragment. After a washing step, samples were incubated for 30 minutes, at room temperature, with Alexa flour 488-conjugated goat anti mouse IgG secondary antibody (Life Technologies). All washing steps and antibodies dilutions were performed using 1 % (w/v) BSA in PBS. Labeled bacteria were washed and fixed for 1 hour, at room temperature, using 2% (v/v) formaldehyde (Carlo Erba Reagents) in PBS. Samples were analyzed with BD FACS Canto™ II system (BD Bioscience) using FlowJo software.

## Results

### NHBA is cleaved by human saliva

Lactoferrin is known to be present in various biological fluids, such as blood, milk, tears, nasal secretions and saliva **[9]**, and it has been reported to be a protease responsible for NHBA cleavage **[7]**. To explore the possibility that NHBA protein may be processed during initial *Neisseria* colonization of the nasopharynx, we assessed whether saliva, which contains hLF, was able to proteoticallly cleave the protein. To this end, human saliva from a pool of donors was incubated with recombinant NHBA (from NZ98/254 strain) protein. After 4 hours, samples were recovered and analyzed by SDS-PAGE. As shown in Fig 1, saliva treatment of NHBA protein gave rise to two main fragments with an apparent molecular weight of approximately 40 kDa and 20 kDa (lane 6), similarly to those produced by hLF **[7]**. Significantly, a recombinant NHBA mutant protein, wherein all Arginines of the Arginine rich region have been deleted,deleted (NHBA ΔArg), was not cleaved in saliva (lanes 9 and 10), indicating that the presence of the Arginine rich region was necessary for the proteolytic cleavage.

**Fig. 1.**
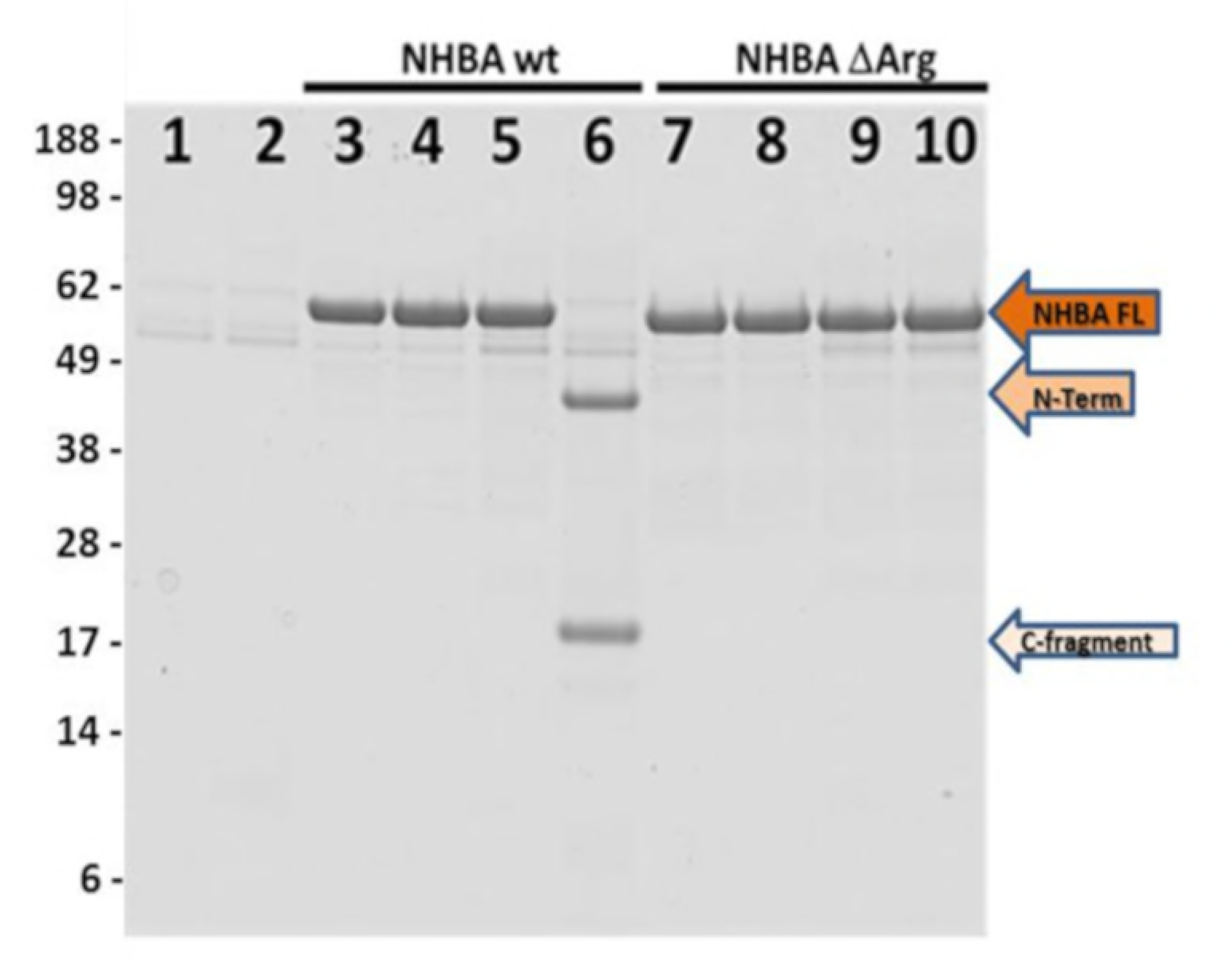
Saliva promotes NHBA cleavage. SDS PAGE of NHBA incubated with human saliva. Lane 1: pool of saliva; lane 2: pool of saliva incubated for 4 hours at 37°C; lane 3: NHBA wt recombinant protein alone; lane 4: NHBA wt recombinant protein incubated for 4 hours at 37°C; lane 5: NHBA wt recombinant protein and saliva; lane 6: NHBA wt recombinant protein incubated with saliva 4 hours at 37°C; lane 7 : NHBA ΔArg recombinant protein alone; lane 8: NHBA ΔArg recombinant protein incubated for 4 hours at 37°C; lane 9 : NHBA ΔArg recombinant protein with saliva; lane 10: NHBA ΔArg recombinant protein incubated with saliva for 4hours at 37°C.

### Lactoferrin is not the main actor of NHBA cleavage in human saliva

To verify whether the protease responsible for NHBA cleavage in saliva was the hLF, proteins contained in saliva were partially fractionated by Anion Exchange Chromatography. Proteins were eluted by increasing ionic strength (Fig. S1), and each recovered fraction was examined for both the ability to cleave NHBA protein and for the presence of hLf. The proteolytic activity of each fraction on NHBA protein was monitored by SDS-PAGE analysis (Fig. 2B), while the presence of hLf was assessed by Western blot analysis (Fig. 2A). Surprisingly, while the hLF was mainly detected in fractionsfrom 33 to 37, from 48 to 50 and 59 and 60, the majority of the proteolytic cleavage of NHBA was observed from fraction 50 to fraction 56, suggesting that hLF was not the main factor, in saliva, responsible for cleavage of NHBA. To further confirm this experimental evidence, either the fractions with major cleavage activity (52-54) or those containing hLF (33-37) were pooled and concentrated, and their ability to cleave NHBA was tested. As shown in Fig 3, a pool of fraction 52-54 completely processed NHBA protein after overnight incubation (lane 4). On the contrary, only a minimal activity was visible for a pool of fractions 33-37 positive for hLf (lane 5). These experimental evidences led to the conclusion that, in addition to hLF, another unknown human protease in saliva was able to cleave NHBA protein and cleavage of NHBA in saliva resulted mainly from its activity.

**Fig. 2.**
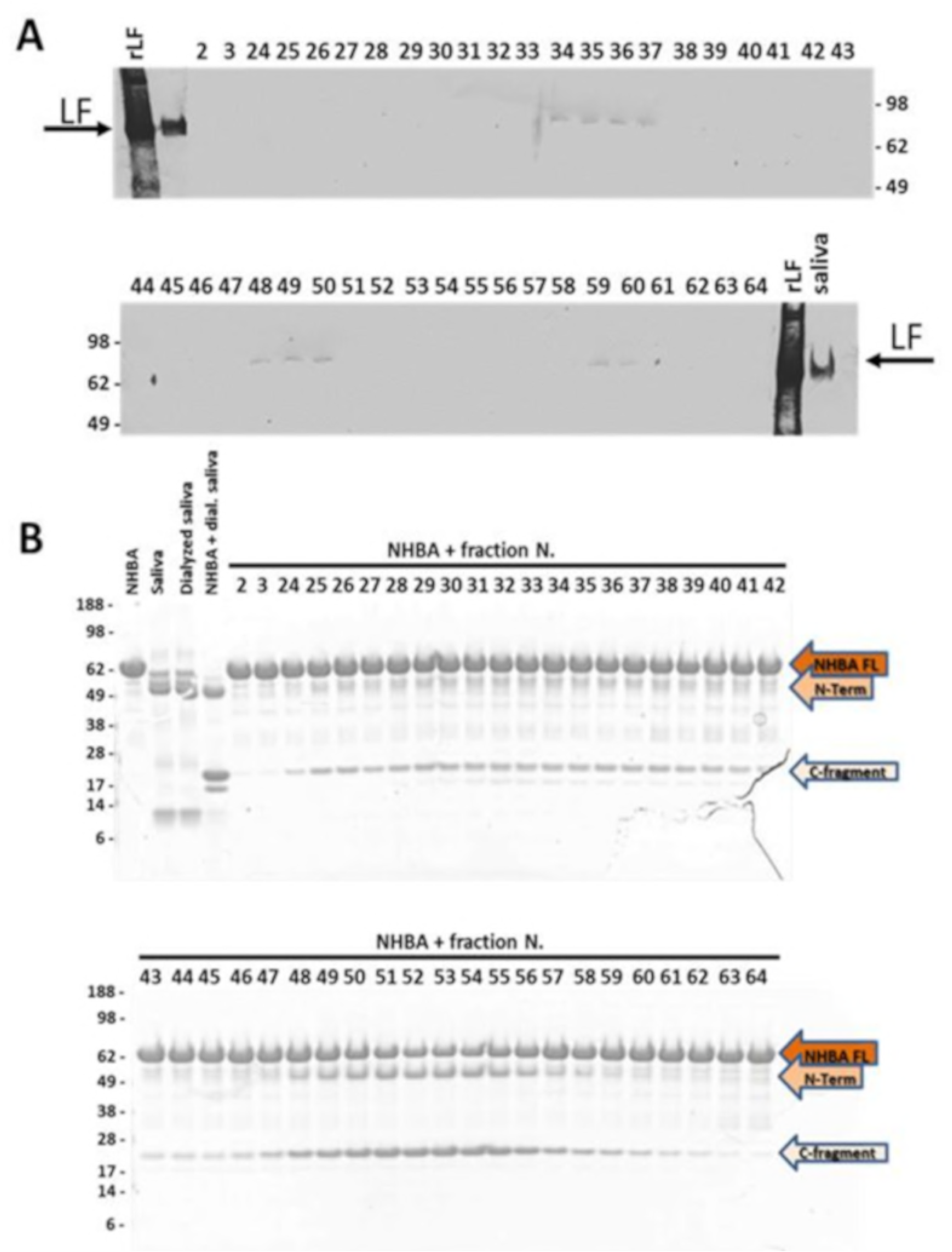
Lactoferrin in Saliva fractions and their activity on NHBA cleavage. Western Blot on saliva recovered fractions against lactoferrin. Recombinant lactoferrin (rLF), saliva (saliva) and the fractions from AEC were loaded on the gel. Lactoferrin is present in frations from 34-37, 48-50 and 59-60. B) SDS Page of the fractions obtained through the saliva partial purification incubated with NHBA. All the samples were maintained at 37°C ON and then loaded on SDS-Page. In the first four lanes, NHBA recombinant protein, saliva, saliva post dialysis and saliva post dialysis incubated with NHBA as reported as control. From number 2 to number 64, are reported the different fractions obtained from saliva purification after incubation with NHBA recombinant protein.

**Fig. 3.**
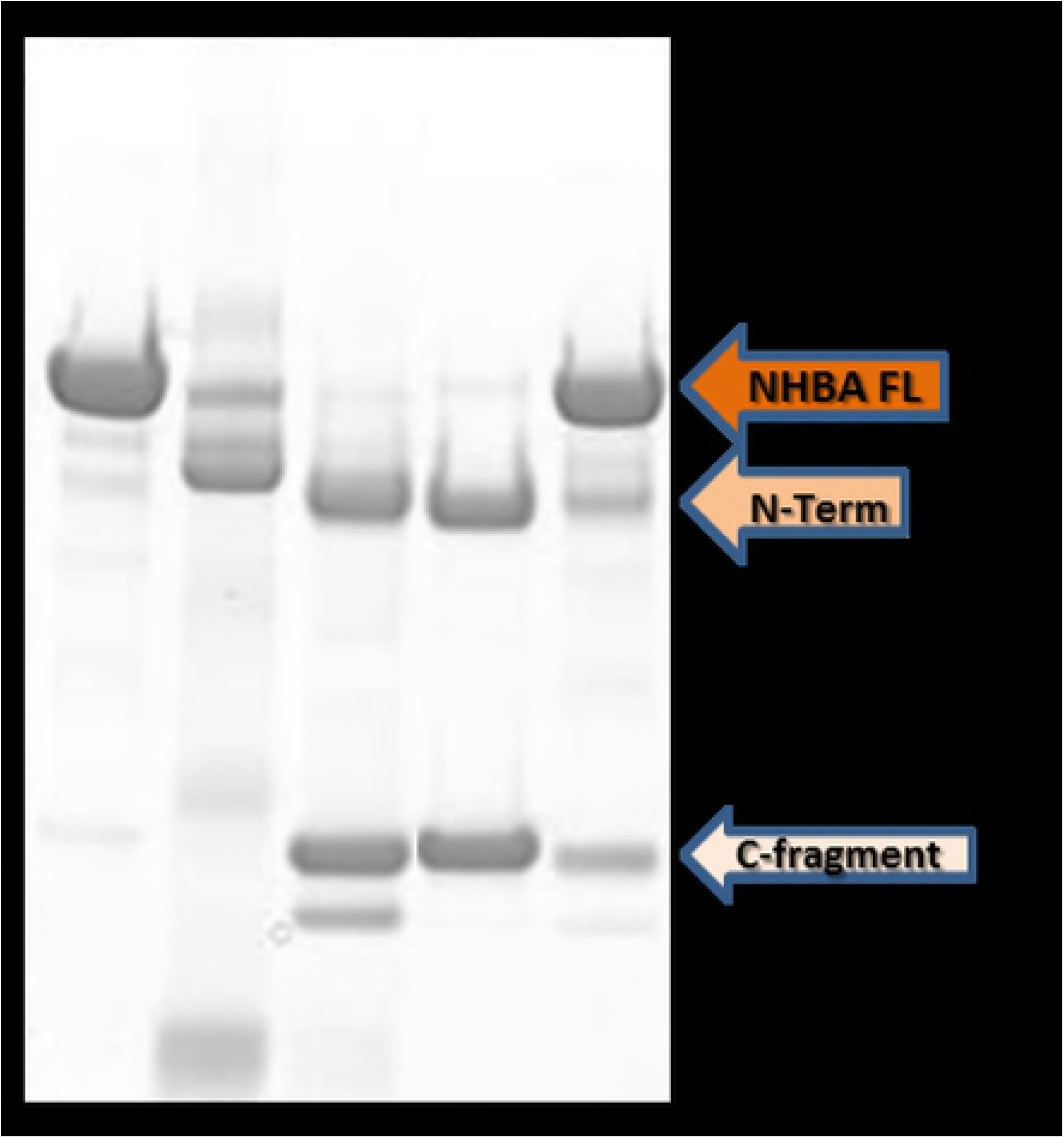
Lactoferrin in saliva is not the main responsible for NHBA cleavage. SDS Page on NHBA processed by recovered and concentrated saliva fractions. All the samples were maintained at 37°C ON and then loaded on SDS-Page. Lane 1: NHBA recombinant protein; lane 2: saliva; lane 3: NHBA recombinant protein incubated with saliva; lane 4: NHBA recombinant protein incubated with pool of fractions from 52-54 concentrated; lane 5: NHBA recombinant protein incubated with concentrated pool of fractions positive for lactoferrin in Western Blotting.

### Kallikrein 1 is the main factor responsible for NHBA cleavage in human saliva

To identify the main protease responsible for NHBA processing in saliva, a pool of fractions (52-54) with the highest NHBA proteolytic activity was analyzed by mass spectrometry based proteomics. Proteins were therefore precipitated, digested with trypsin and the resulting peptides were subjected to LC-MS/MS analysis. Protein identification from mass spectrometry data was performed against the *Homo sapiens* database publically available in NCBInr. Among the identified proteins, we found the protease kallikrein 1 (hKLK1), whose identity was assigned by the MS/MS sequence of two peptides (Fig. 4). This isoform of kallikrein is specifically expressed in human kidney, pancreas and salivary glands **[21]** and its cleavage site is located downstream a positively charged amino acid (such as Lysine or Arginine). Hence, it could be a plausible candidate for cleavage of NHBA in saliva, since NHBA contains an Arginine-rich region that is highly conserved among different *N. meningitidis* strains and that could act as target for hKLK1 proteolytic cleavage. To further investigate this matter, recombinant human kallikrein1, kallikrein1 purified from human plasma (commercially available) and human saliva were incubated with recombinant NHBA protein. SDS-PAGE analysis showed that both recombinant hKLK1 (lane 3-6, Fig 5) and hKLK1 purified from human plasma (lane 7-10, Fig. 5) were able to proteoliticcaly cleave NHBA with fragmentation patterns identical to that induced in saliva (lane 2, Fig. 5). Proteolytic cleavage resulted in the generation of two main fragments (highlighted with red arrow in Fig. 5), which amount increased, increasing the amount of kallykrein, in a dose-dependent manner. Moreover, the NHBA fragments generated by the cleavage showed an apparent molecular weight comparable to the fragments produced by hLf cleavage. **[7]** These results indicated that KLK1 is the main protease responsible for NHBA processing in saliva.

**Fig. 4.**
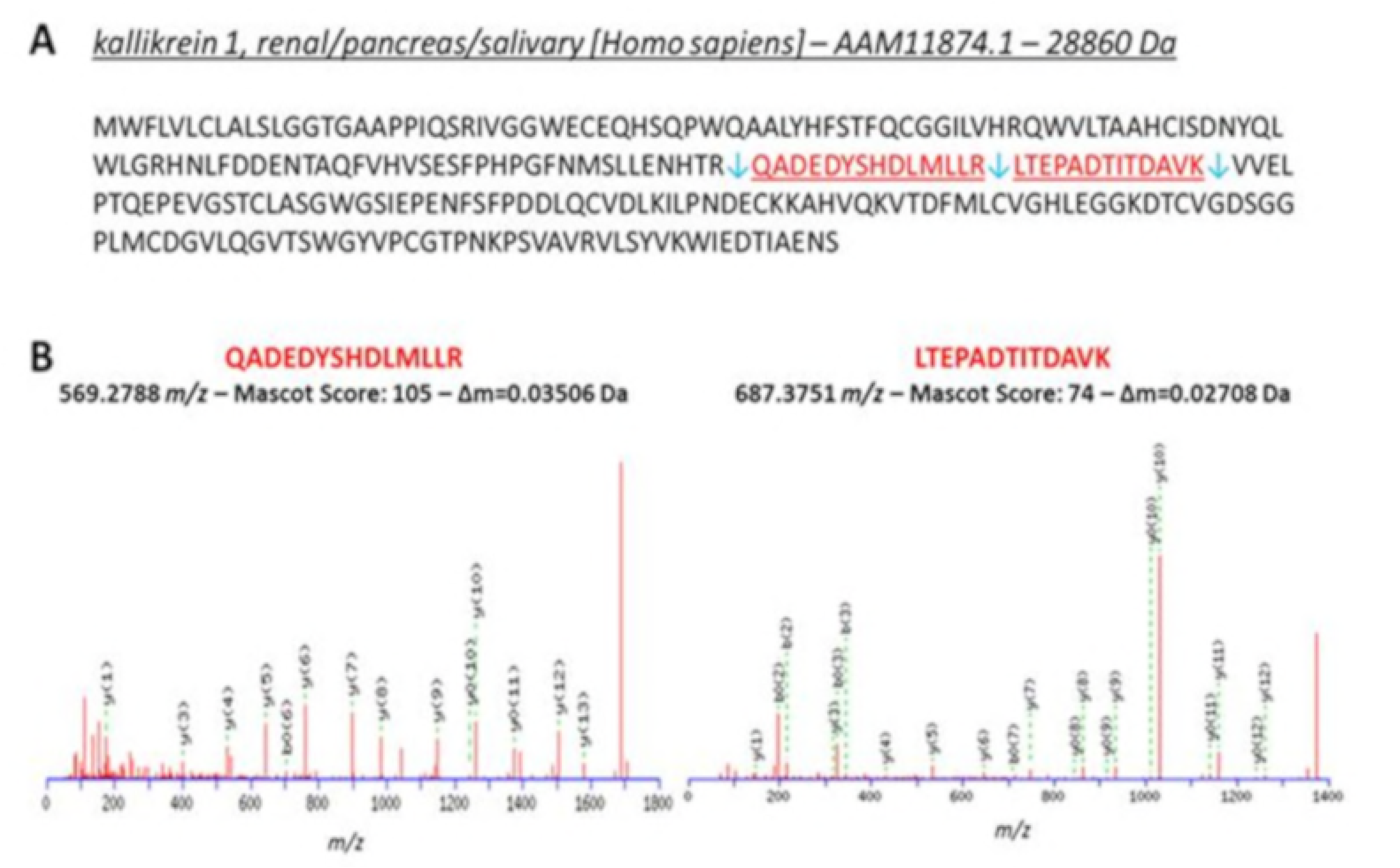
The protease kallikrein 1 was identified in NHBA-cleaving fractions from human saliva. **(A)**Primary sequence of kallikrein 1 identified from saliva fractions able to cleave NHBA. Identified peptides by LC-MS/MS are highlighed in red. Blue arrows indicate trypsin cleavege sites. **(B)** MS/MS spectra of the peptides identified from kallikrein: m/z values, Mascot score and experimental delta mass are reported.

**Fig. 5.**
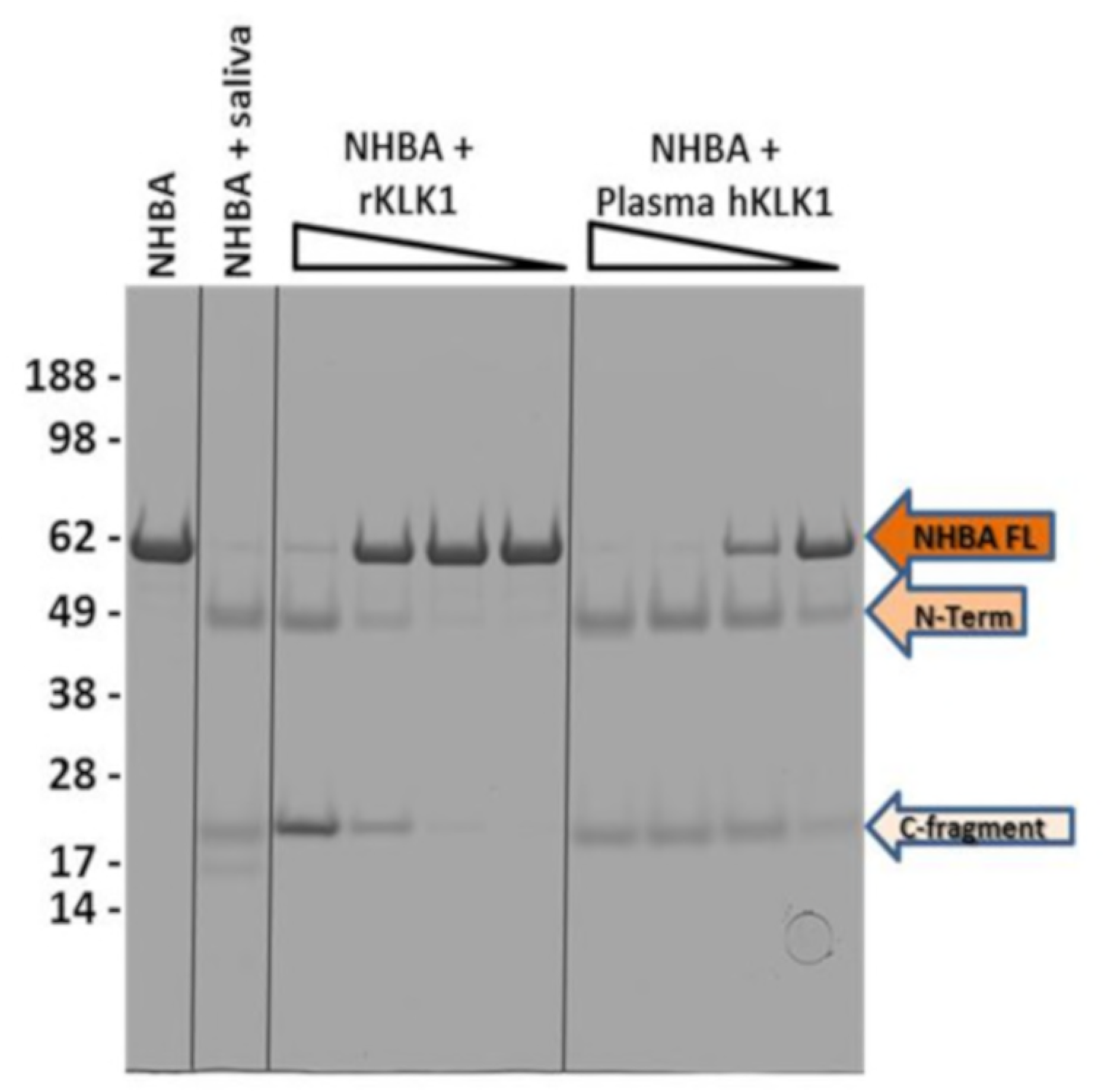
Kallikrein processes NHBA in-vitro. SDS-PAGE of NHBA cleavage by human kallikrein. All the samples were maintained at 37°C ON and then loaded on SDS-Page. Lane 1: NHBA alone; Lane 2: NHBA incubated with human saliva; Lane 3-6 : NHBA incubated with recombinant KLK1 (rKLK1) at dilution from 1:500, 1:5000, 1:50000 and 1:500000; Lane 7-10 : NHBA incubated with human KLK1 (hKLK1) purified from human plasma at dilution from 1:500, 1:5000, 1:50000 and 1:500000.

### Kallikrein and human lactoferrin promote the formation of two identical fragments

To identify the exact cleavage site within the NHBA protein promoted by hKLK1 and saliva, we purified the C-terminal fragments generated by their cleavage. Therefore, NHBA was incubated with saliva or with rhKLK1 (recombinant human kallikrein 1) overnight and the resulting fragments were separated by affinity chromatography. Since NHBA was His-tagged at the C-terminus, we used NiNtA resin to bind and purify the cleaved C-terminal fragments, while N-terminal fragments were not bound to the column and released in the flow through (Fig. 2S).

Once the cleaved C-term fragments generated by human saliva and recombinant human kallikrein1 were purified, they were analyzed by mass spectrometry to determine their precise molecular weight, thus allowing identification of the cleavage site of hKLK1 and saliva within the NHBA protein. Intact mass determination was performed by positive electrospray ionization using a Q-TOF mass spectrometer. For the NHBA C-fragment derived from saliva cleavage, we detected a molecular weight of 20246.57 ± 0.14 Da that was compatible with the primary sequence of the C-terminal region of NHBA starting from Ser^288^ (Fig. 6A). Similarly, the NHBA fragment derived from recombinant human kallikrein1 cleavage showed a molecular weight of 20247.3 Da ± 2 Da (Fig. 6B) that was also compatible with the same C-terminal sequence starting from Ser^288^. We therefore concluded that the cleavage site of both hKLK1 and saliva was located immediately downstream to the Arginine rich region at the level of Ser 288, and corresponded to the cleavage site of hLF **[7]**.

**Fig. 6.**
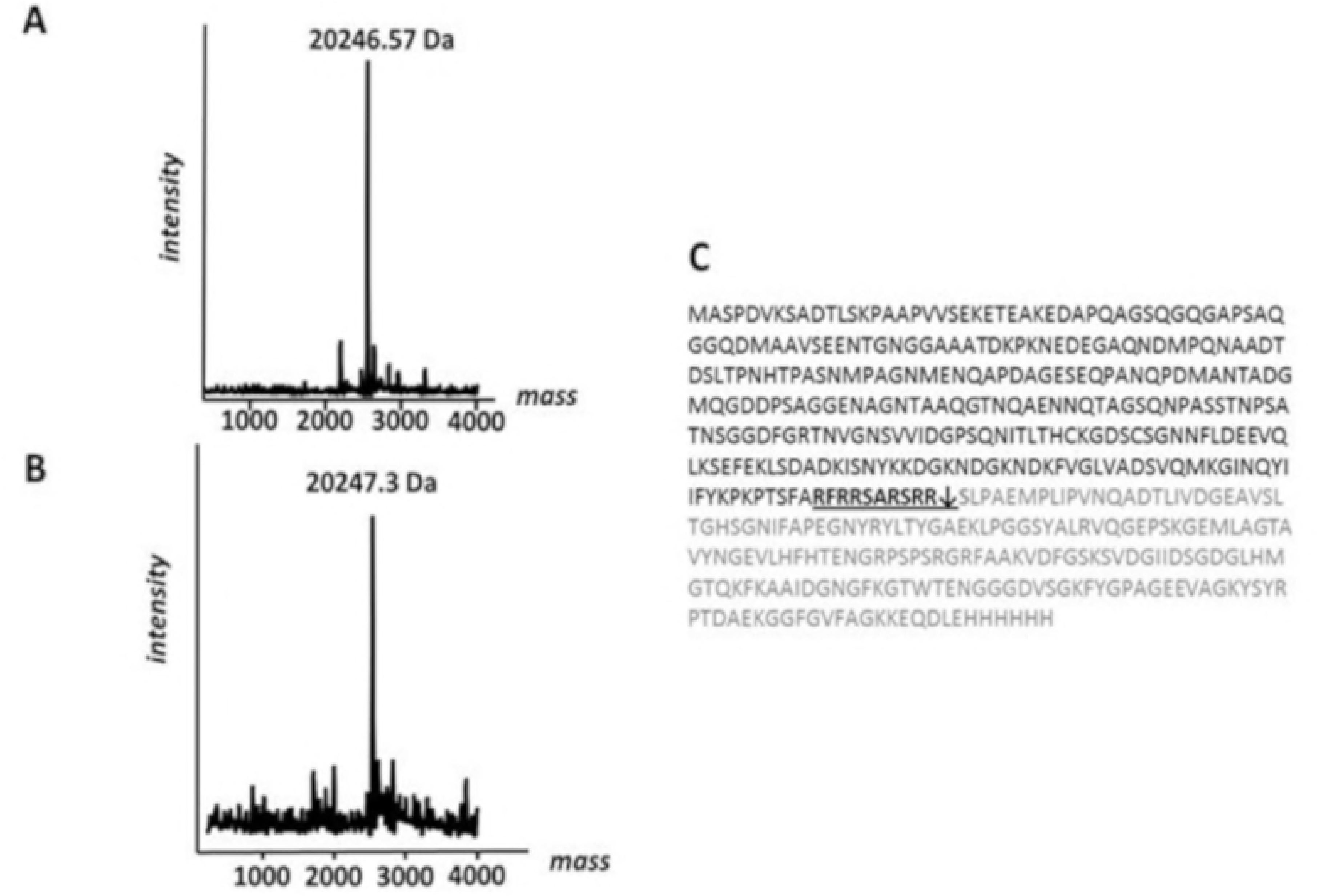
Intact mass measurement of NHBA C-terminal fragments obtained from kallikrein cleavage. Intact mass determination was performed by ESI-Q-TOF. A) The NHBA C-fragment after saliva cleavage showed an observed molecular weights of 20246.57 Da, compatible with the primary sequence of C-terminal region of NHBA starting from Ser288. B) NHBA fragment after recombinant human kallikrein1 cleavage showed a molecular weight of 20247.3 Da. C) Amino acid sequence of recombinant NHBA (NZ) full-length protein: Arg-rich motif is indicated in bold, cleavage site of epithelial cells proteases is indicated with a black arrow and amino acid sequence of newly generated C-terminal fragment is highlighted in gray.

### NHBA and kallikrein in *Neisseria meningitidis* bacteria

Since *in-vitro* data proved that hKLK1 was the main protease in saliva responsible for the proteolytic cleavage of NHBA into two main fragments, the N-and C-terminal, we further verified their ability to process NHBA on native protein expressed on the surface of live bacteria.

To test NHBA processing on live bacteria, *Neisseria meningitidis* MC58 wild-type strain and the relative knock-outs for *nalP gene* (coding for the meningococcal serine protease able to cleave NHBA) and *NHBA* gene, as a control strains, were grown at 32°C to simulate the nasopharyngeal environment **[14],** and then treated with hKLK1 or saliva. After 30 minutes, samples were collected and the presence of the NHBA C-terminal domain was assessed either into supernatants by Western blot analysis (Fig. 7A) or on bacterial surface by FACS analysis (Fig. 7B). As previously reported **[7]**, the NHBA-C terminal fragment derived for cleavage of meningococcal NalP protease, named C2, could be detected in supernatant of MC58 wt (Fig. 7A, lane 4). In addition, a shorter NHBA C-terminal fragment was observed in the supernatant of bacteria incubated with hKLK1 or saliva (Fig. 7A, lane 5 and 6) with an apparent molecular weight identical to the NHBA C-terminal fragment generated by hLF, named C1 **[7]**. Moreover, while NHBA protein was not cleaved by meningococcal proteases, as in MC58Δ*nalp*, the NHBA C-terminal fragment was only detected in the supernatant of bacteria incubated with hLKL1 or saliva (Fig.7A, lane 8 and 9), indicating that both treatment led to an accumulation of NHBA C-terminal fragment into the supernatant. Of note, we observed the NHBA full-length protein in all samples of both MC58 wt (lanes 4-6) and MC58Δ*nalp* strain (lanes 7-9), while it was not present in the MC58 Δ*NHBA* strain (lanes 10-12). All samples treated with saliva showed a signal at a molecular weight a little bit higher than the C-fragment (red arrow in lane 6, 9 and 12), which is an unspecific signal of the polyclonal serum used on the saliva sample.

**Fig. 7.**
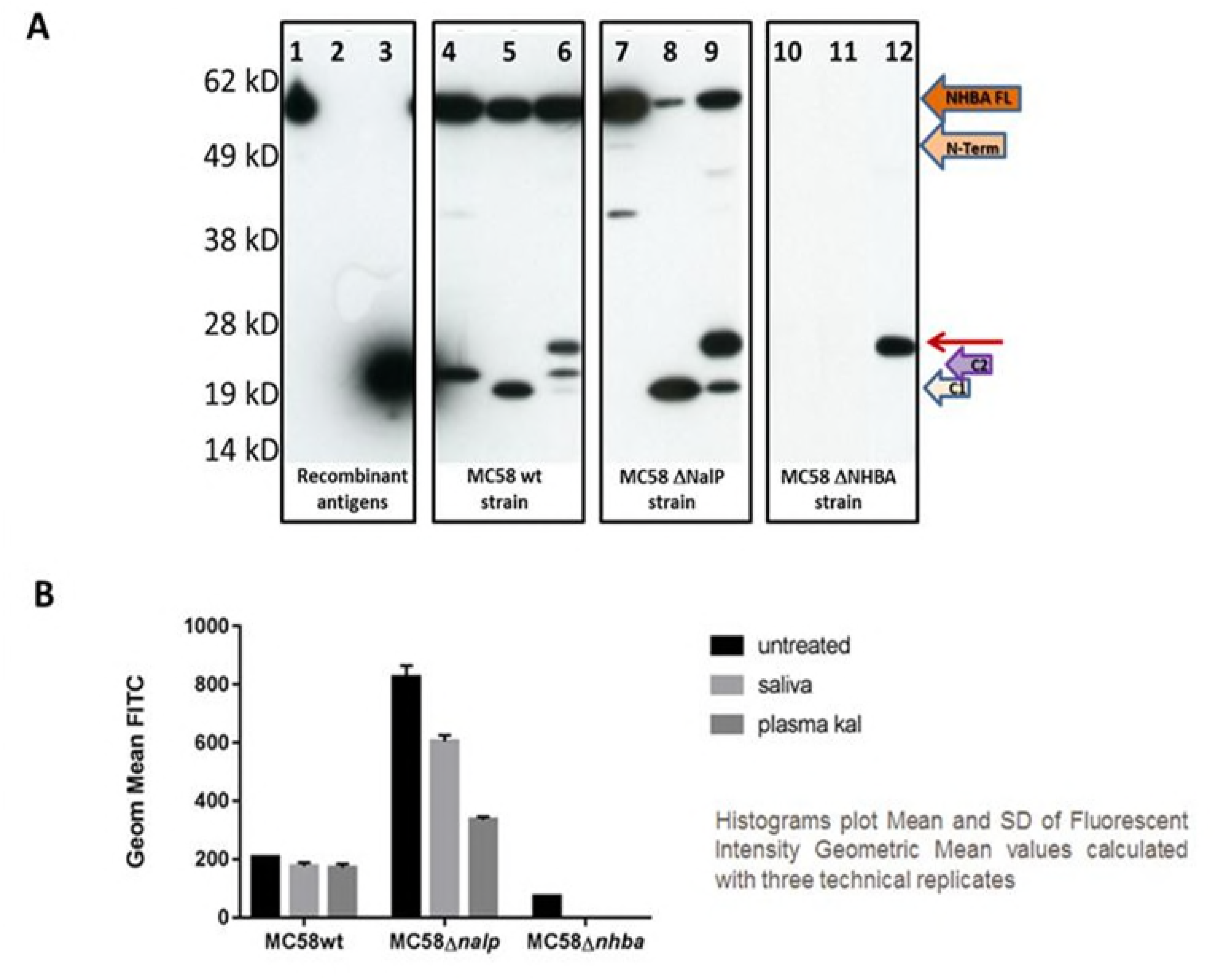
NHBA cleavage by human kallikrein and saliva occurs on live bacteria. Western Blotting assay, using α-NHBA - C - fragment polyclonal antibody. Lane 1: NHBA wild type recombinant protein; lane 2 : NHBA - AB recombinant fragment; lane 3 : NHBA recombinant C-fragment; lane 4: MC 58 wild type supernatant; lane 5: MC58 wild type supernatant following incubation with kallikrein; lane 6: MC58 wild type supernatant following incubation with saliva; lane 7: MC 58 ΔNalP supernatant; lane 8: MC58 ΔNalP supernatant following incubation with kallikrein; lane 9: MC58 ΔNalP supernatant following incubation with saliva; lane 10: MC58 ΔNHBA supernatant; lane 11: ΔNHBA supernatant incubated with kallikrein; lane 12: MC58 ΔNalP supernatant incubated with saliva. B) FACS analysis of MC58wt, MC58Δnalp and MC58Δnhba strains stained with anti NHBA C-terminal-FITC antibody. Bacteria were incubated with for 30 minutes with human kallikrein, saliva or with medium, as negative control. Histograms plot the geometric mean of Fluorescence Intensity (FITC). Error bars denote standard deviation.

These data show that NHBA C-fragments produced by either hKLK1 or saliva have the identical molecular weights, whereas the molecular weight of the C-fragment produced by NalP cleavage (C2 fragment) is different.

Evidence that hKLK1 and saliva cleaved NHBA protein on live bacteria was further supported by FACS analysis. Therefore, following protease treatments, bacteria were stained with anti NHBA C-terminal – FITC antibody and the amount of C-terminal domain on surface-exposed NHBA protein was determined by measuring fluorescent intensity. As reported in Fig. 7B, no differences were observed in MC58wt strain comparing treated and untreated bacteria. In addition, the basal level of NHBA C-terminal domain present on the bacterial surface was very low in untreated samples. Likely, most of the NHBA exposed on the bacterial surface was already processed by meningococcal proteases at the time of treatment, since this strain highly expressed Nalp protease. On the contrary, for the *nalp* knockout mutant, in which most of the surface exposed NHBA protein was present in its full-length form, the amount of NHBA C-terminal domain on the bacterial surface decreased after treatment with hKLK1 or saliva. This reduction was particularly significant for samples incubated with hKLK1.

These results indicate that both saliva and hKLK1 were able to cleave NHBA on live bacteria and the generation and release of NHBA C-terminal domain from the bacterial surface resulted in the accumulation of C-terminal fragment into the supernatant.

## Discussion

The role of NHBA in *Neisseria meningiditis* pathogenesis is still unclear. Previous studies have shown that NHBA is able to bind Heparin and GAGs on epithelial cells through the Arg-rich region **[16]**, supporting the hypothesis that NHBA can not only enhance bacterial survival in human serum, but can also play a crucial role during the adhesion step in *Neisseria meninigitidis* host colonization **[15].** NHBA is cleaved by two specific proteases, NalP, a surface-exposed *Neisseria* serine protease, and human lactoferrin (hLf), present in several human biological fluids like serum, milk and saliva **[7]**. The fragment generated by proteolytic cleavage can induce a toxic effect on endothelial cells by altering the endothelial permeability **[15].** In this study, we have assessed whether the proteolytic cleavage of NHBA could happen in saliva, being the nasopharynx the only known *N. meningitidis* bacterial reservoir, and the first tissue where the bacterium starts its colonization. Understanding the mechanism which allows the bacterium to pass through the human nasopharyngeal epithelium and start its invasive phase is a crucial step to develop a preventive treatment for meningitis disease. For this purpose, we evaluated the possibility that NHBA could be processed by human saliva, the main fluid present in the nasopharyngeal tract. In this work, we demonstrate that NHBA is cleaved *in vitro* by human saliva into two main fragments, an N-terminal fragment (AB) and a C-terminal fragment (C1), and that C1 is the same fragment generated by hLf processing of NHBA; we found, however, that the main actor for NHBA processing in saliva is not hLf but kallikrein1 (KLK1). KLK1 is a human serine protease that cleaves one aminoacid downstream of positively charged amino-acids and is present in several biological fluid as plasma, lymph, urine, saliva and pancreatic juice. We demonstrated that hKLK1 cleavage site on NHBA happens on the Ser 288 just downstream the Arg-rich region, confirming that hKLK1 processes NHBA at the same identical site previously reported for hLf **[7].**

We have so far three proteases able to cleave NHBA *in vitro,* upstream or downstream the Arg-rich region: meningococcal NalP protease (upstream), that generates fragment C2 containing the Arg-rich domain, and hLf and hKLK1 (downstream), two human proteases generating fragment C1, in which the Arginine-rich domain is absent. Both fragments are released from the bacteria in to the environment **[15].**

To further characterize and understand the possible biological relevance of NHBA processing we compared the activity of human saliva with the newly identified NHBA cleaving protease, hKLK1, on the bacterial surface by testing their ability to process NHBA on live bacteria. We showed that NHBA is processed in ΔNalP MC58 strain by saliva and by hKLK1 generating an identical peptide with a MW equal to the C1 fragmen,t and it is also processed by NalP, generating the C2 fragment. Both fragments, C1 and C2 are released in the culture medium.

We demonstrated that NHBA can be cleaved on live bacteria releasing two main fragments, C1 and C2, from the bacterial surface. Hence, we speculated whether the two different C-term fragments could play a different role in NHBA mediated *Neisseria* pathogenesis.

The important role of the NHBA Arg-rich region for host colonization and invasion was demonstrated previously for Heparin binding **[7]** and *Neisseria meningitidis* adhesion to Human Ephitelial Cells **[16].**

It was also shown that C2, but not C1, increases endothelial permeability with a mechanism involving the production of oxygen radicals and the phosphorylation and degradation of adherents-junction protein VE-cadherin. **[15]**. Since the C2 fragment contains the Arg-rich domain it is plausible that this fragment might interact with secreted or cell-associated proteoglycans in order to exert its biological role **[15].** Moreover, the C2 fragment seems to be critical for invasion of human tissues, hence for virulence, and different studies showed the possibility that it is naturally released in the bacterial environment. We can also assume that the C1 fragment, released by *N. meningitidis* in the nasopharyngeal tract in its first colonization phase and processed *in vivo* by KLK1 present in saliva and therefore lacking the Arginine-rich domain responsible for binding to heparin, is not able to adhere on endothelial cell proteoglycans and enter the blood stream **[15].** Therefore, hKLK1, and possibly to a lesser extent hLf, producing an inactive C-fragment, may have a protective role for the host.

The Arg-rich region is highly conserved among different NHBA proteins and across MenB strains **[7]**, suggesting a crucial role in NHBA function in its ability to bind heparin. However, there is yet no concrete explanation why, in the pharyngeal tract, three proteases are involved in NHBA processing, with meningococcal NalP targeting the Arg-rich region upstream, and human KLK1 and human lactoferrin targeting it and downstream. Also it is not clear why a domain which is the target of the front-line host defense mechanism is so widely conserved among *Neisseria* subtypes. Bacterial colonization in the nasopharyngeal tract is the prerequisite for the *N. meningitidis* invasive phase. Successful colonizers must attach to the epithelial lining, grow on the mucosal surface, evade the host immune response, and penetrate in to the bloodstream. However, the invasive behavior is not part of the normal meningococcal life cycle since, once the colonizer have entered the bloodstream or the central nervous system, they cannot be easily transmitted to other hosts, and blood not only constitutes an immunologically challenging compartment but also an oxygen-limiting environment **[23]**. Environmental conditions which force the switch of *N. meningitidis* from carriage to invasive are not yet known, and the prevailing way of NHBA cleavage, by NalP producing C2 active fragment or by KLK1 producing C1 inactive fragment, may play a role in signaling to the bacteria the environmental changes required for initiation of the invasive phase. This aspect opens the possibility that NHBA processing by host proteases would not be solely a host defense mechanism, but could be crucial for the bacterium decision-making process in the carriage-to-virulence switch, hence its conservation among *N. meningitidis* subtypes.

Further studies may consider investigating the environmental conditions which activate or inhibit NalP or KLK1 functions, and how these could affect meningococcus virulence and invasiveness.

## Acknowledgments

The authors want to thank Silvana Savino, Mariagrazia Pizza, Beatrice Aricò and Vega Masignani (GSK, Siena, Italy) for useful discussion, and Antonietta Maiorino for rewing the paper in the original language.

Bexsero is a trademark of the GSK group of companies.

## Authors’ contribution

Conceived and designed the experiments: Elena Cartocci

Performed *in vitro* experiments: Elisa Pantano

Performed depletion of lactoferrin from saliva: Sara Marchi, Vincenzo Nardi-Dei

Performed FACS analysis: Silvia Rossi Paccani and Martina Di Fede

Performed MS analysis: Massimiliano Biagini

Analyzed the data: Elisa Pantano, Elena Cartocci, Sara Marchi, Vincenzo Nardi-Dei, Massimiliano Biagini

Wrote the paper: Elisa Pantano

Reviewed and approved the final version of the manuscript: all authors.

## Supporting information

**S1 Fig**. **Saliva fractionation on AEC.**

Chromatogram of the Anion Exchange chromatography. Red box highlights fractions with major protease activity on NHBA.

**S2 Fig**. **NHBA fragments purification after digestion with saliva and kallikrein.**

A)SDS PAGE of C-fragment purification after saliva NHBA processing. Lane 1: NHBA recombinant protein alone; lane 2: NHBA recombinant protein incubated with saliva ON at 37°C; lane 3: NiNtA flow thorught fractions; lane 4 and 5: NiNtA column washes; lane 6 and 7: elution fractions from the NiNtA column which contain the C fragment. B) SDS PAGE of C-fragment purification from recombinant kallikrein NHBA processing. Lane 1: NHBA recombinant protein alone; lane 2: NHBA recombinant protein incubated with recombinant kallikrein ON at 37°C; lane 3 : NiNtA flow thorught fractions; lane 4 and 5: NiNtA column washes; lane 6 and 7 : elution fractions from the NiNtA column which contain the C fragment.

